# The endoplasmic reticulum associated degradation adaptor Sel1L regulates T cell homeostasis and function

**DOI:** 10.1101/2021.05.22.445275

**Authors:** Alexander T. Dils, Luis O. Correa, J. Paige Gronevelt, Lu Liu, Padma Kadiyala, Qing Li, Shannon A. Carty

**Affiliations:** Division of Hematology-Oncology, Department of Internal Medicine, Ann Arbor, MI 48109; Immunology Graduate Program, Ann Arbor, MI 48109; Rogel Cancer Center, University of Michigan, Ann Arbor, MI 48109

**Keywords:** Sel1L, ERAD, mTOR, T cell, quiescence, CD8

## Abstract

Suppressor/Enhancer of Lin-12-like (Sel1L) is a critical adaptor for endoplasmic reticulum-associated degradation (ERAD), a process that maintains cellular protein quality control through degradation of misfolded proteins. Here we investigate the role of Sel1L in T cell homeostasis and function. T cell-specific deletion of Sel1L profoundly impairs peripheral T cell survival and promotes apoptotic cell death. Furthermore, Sel1L is required to maintain naïve CD8^+^ T cell homeostasis in a cell-intrinsic manner with loss of quiescence as evidenced by increased proliferation. Sel1L-deficient T cells exhibit enhanced activation of the mammalian target of rapamycin (mTOR) pathway and altered cellular metabolism, including increased cellular reactive oxygen species, mitochondrial mass and mitochondrial membrane potential in the naïve CD8^+^ T cell compartment. Furthermore, loss of Sel1L impaired CD8^+^ T cell immune responses following bacterial infection. These results demonstrate a novel role for Sel1L/ERAD in T cell homeostasis and function.

## Introduction

For cells to maintain cellular health and appropriate function, there must be tight control of protein synthesis, processing/folding and degradation processes. Protein homeostasis (proteostasis) is maintained partially through the ER-associated degradation (ERAD) pathways. ERAD complexes contain ER transmembrane proteins that recognize misfolded proteins in the ER and translocate them to the cytosol for ubiquitination and subsequent proteasomal degradation (Qi et al., 2017). There is a constant demand for ERAD-mediated surveillance to clear aberrant proteins from the ER, which occurs when a native structure fails to form due to mutation, translational mis-incorporation or stochastic inefficiency in adopting native conformation or forming protein complexes. Sel1L acts as an important adaptor of the ERAD complex, recognizing misfolded proteins in the ER and recruiting them to be exported to cytosol for proteasomal degradation. Sel1L also binds to and stabilizes hydroxymethylglutaryl reductase degradation protein 1 (Hrd1), the E3 ubiquitin ligase of the ERAD complex, thus making it essential for Hrd1-ERAD function (Sun et al., 2014).

There is emerging data about the importance of protein homeostasis for T cell fate and function in settings of cellular stress, such as during infection, inflammation, and the tumor microenvironment. ERAD through Hrd1 is required for proliferation, IL-2 production and T_H_1/T_H_17 differentiation following TCR activation in murine CD4^+^ T cells (Kong et al., 2016) and is required for regulatory T cell (T_reg_) stability under inflammatory conditions (Xu et al., 2019). However, little is known about Sel1L’s role in T cells.

Here we report that Sel1L is required for peripheral T cell survival and maintenance of naïve CD8^+^ T cell quiescence. Loss of Sel1L in later stages of T cell development results in largely normal thymocyte numbers with a minor decrement of CD4 single-positive (SP) and CD8SP thymocytes but results in a marked decrease in peripheral T cells compared to littermate controls. Sel1L protects against apoptotic cell death and promotes peripheral T cell survival in a cell-intrinsic manner. In addition, Sel1L is required to maintain naïve CD8^+^ T cell homeostasis and quiescence. Sel1L loss leads to mTORC1 activation and altered cell metabolism in T cells. Furthermore, Sel1L is required for optimal CD8^+^ T cell immune responses to *Listeria monocytogenes* infection. Together, these findings identify a key role for Sel1L/ERAD for T cell survival, homeostasis, and anti-bacterial immunity.

## Results

### Sel1L promotes peripheral T cell survival

To examine the role of Sel1L in mature T cells, we crossed mice with loxP-flanked Sel1L alleles (Sel1L^fl/fl^; (Sun et al., 2014) with mice expressing the Cre recombinase transgene under the control of the *Cd4* promoter (Sel1L^fl/fl^CD4Cre^+^; referred herein as Sel1LKO) to delete the Sel1L starting in the double positive (DP) stage of thymocyte development. Analysis of messenger RNA and protein demonstrated efficient deletion of Sel1L in the T cells (Figure 1A). When misfolded proteins accumulate, they form protein aggresomes (Johnston et al., 1998). In Sel1L-deficient T cells, we noted a higher level of protein aggresomes, as measured by the flow cytometric reagent Proteostat, compared to wild-type cells (Figure 1B), as would be predicted if ERAD was disrupted. Littermate control (wild-type; WT) and Sel1LKO mice had a similar absolute number of total thymocytes, as well as CD4^−^ CD8^−^ double-negative (DN), CD4^+^CD8^+^ double-positive (DP), though had a slight decrease in the absolute number of CD4 single-positive (SP) and CD8SP cells (Figure 1C). In the spleen, Sel1L deletion led to a marked reduction in the frequency and absolute number of T cells affecting both CD4^+^ and CD8^+^ T cells (Figure 1D) without an alteration in the ratio of CD4^+^ to CD8^+^ T cells (data not shown). There was also no difference in the frequency of Foxp3^+^ regulatory T cells (Tregs) amongst the peripheral CD4^+^ T cell population (Supplementary Figure 1).

**Figure 1.**
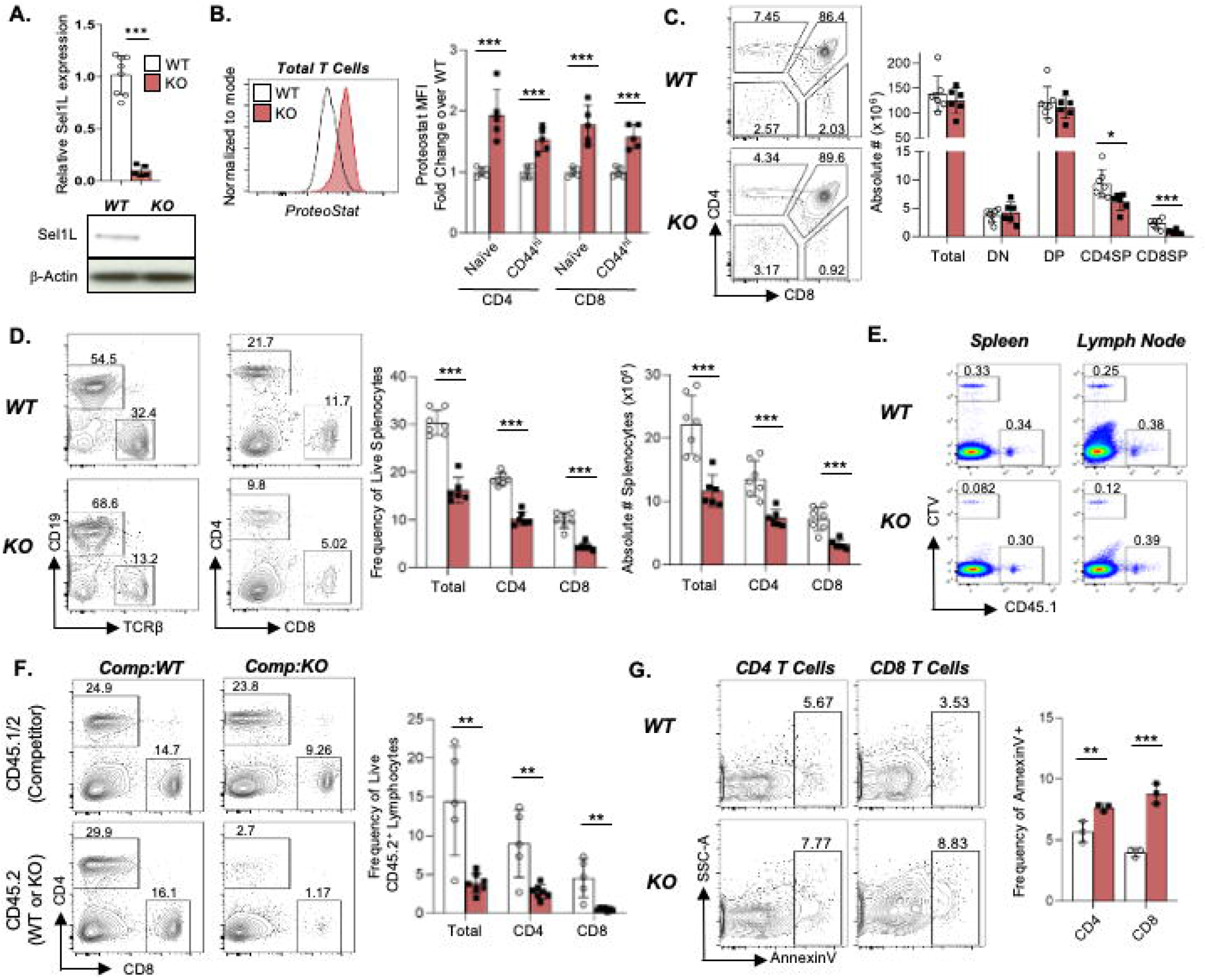
Sel1L maintains peripheral T cell survival. Characterization of thymic and peripheral T cell subsets in Sel1LKO mice and littermate controls (WT), aged 6 10 weeks. A) *Top,* Sel1L expression, relative to b-actin, in cDNA generated from WT and Sel1LKO CD8^+^ T cells; *Bottom,* Sel1L protein expression with b-actin loading control in WT and Sel1LKO CD8^+^ T cells. B) *Left,* Representative flow cytometric analysis of Proteostat in WT and Sel1LKO T cells; *Right,* median fluorescence intensity (MFI) of Proteostat, normalized to WT, in T cell subsets from WT and Sel1LKO mice. C) *Left,* representative flow cytometry of live thymocyte subsets in WT and Sel1LKO, mice. *Right,* frequency and absolute numbers of thymocyte subsets. D) *Left,* representative flow cytometry of TCRb+ splenocytes and CD4^+^, CD8^+^ splenocytes from WT and Sel1LKO mice; *Right,* frequencies and absolute number of peripheral T cell subsets in the spleen of WT and Sel1LKO mice. E) Representative flow plots of splenocytes isolated from CD45.2^+^ hosts which received WT or Sel1LKO T cells (CTV labelled) and CD45.1^+^ ‘spike’ T cells 5 days prior to analysis. F) *Left,* representative flow cytometry of CD4^+^ and CD8^+^ cells among live CD45.2^+^ splenocytes in, mixed BM chimera hosts (CD45.l^+^) which had received 50:50 mixture of CD45.l^+^CD45.2^+^ competitor BM with either WT (CD45.2^+^ CD45.1^-^) or Sel1LKO (CD45.2^+^CD45.1^-^) BM 10-12 weeks prior; *right,* frequency of TCRb^+^ lymphocytes among donor-derived (CD45.2^+^) splenocytes. G) *Left,* Representative flow cytometry of Annexin V staining in CD4^+^ and CD8^+^ T cells in the WT and Sel1LKO mice; *right,* frequency of Annexin V+ population in CD4^+^ and CDS^+^ T cells from the spleens of WT and Sel1LKO mice, nom1alized to WT CD4^+^ T cell population. Data are representative of n=5-8/genotype (A), 5/genotype (B, F), 6-7/genotype (C-E), 8-9/genotype (G), from 2 (B, F) or 3 (A, C-E, G) independent experiments. Data are mean± SD. *p < 0.05, **p <0.01, ***p <0.001 (unpaired t-test). See also Supplementary Figure 1.

We next tested if the reduction of peripheral T cells was due to a survival defect in the setting of Sel1L deletion. A mixture of equal numbers of WT or Sel1LKO donor T cells (labelled with CTV) were adoptively transferred with congenic ‘spike’ T cells into lymphoreplete CD45.2^+^ recipient mice. Five days after transfer, the Sel1LKO T cells were scarcely detectable, while wild-type donor cells were easily detected in the spleen and peripheral LNs (Figure 1E), suggesting that Sel1L is critical for T cell survival in vivo.

To determine if the loss of peripheral T cells was a cell-intrinsic defect, we generated mixed bone marrow (BM) chimeras by transplanting a 50:50 mixture of WT or Sel1LKO T cell-depleted BM cells with wild-type competitor (CD45.1^+^CD45.2^+^) T cell-depleted BM into lethally irradiated congenic mice. Reconstituted populations derived from Sel1LKO BM had fewer CD4^+^ and CD8^+^ T lymphocytes compared to those derived from WT BM (Figure 1F). These data indicate that Sel1L acts in a cell-intrinsic manner to maintain the peripheral T cell pool.

We measured Annexin V to interrogate the mechanism by which Sel1L affects peripheral T lymphocyte survival. There was increased Annexin V staining in both Sel1LKO CD4^+^ and CD8^+^ T cells compared to WT T cells (Figure 1G). These data suggest that Sel1L leads to increased apoptosis at steady-state.

Since interleukin (IL)-7 is known to regulate peripheral T cell survival in vivo (Rathmell et al., 2001), we examined IL-7 receptor (IL-7Ra/CD127) expression in WT and Sel1LKO peripheral T cells. We found increased expression of CD127 on both CD4^+^ and CD8^+^ T cells (Supplementary Figure 2A) in the spleen of Sel1LKO mice. Furthermore, to test if Sel1LKO T cells respond to IL-7 survival signals, we incubated WT or Sel1LKO splenocytes in the absence and presence of IL-7 ex vivo for 18hrs and examined survival. WT and Sel1LKO T cells had similar frequency of cell survival in the presence of IL-7 (Supplementary Figure 2B), suggesting that Sel1L loss does not alter IL-7 survival signaling in T cells. IL-7 promotes T cell survival in part through upregulation of the anti-apoptotic factor Bcl2 (Jacobs et al., 2010). We examined Bcl2 expression in WT and Sel1LKO T cells and consistently found increased Bcl2 expression in the Sel1LKO cells (Supplementary Figure 2C). Together these data indicate that Sel1L does not regulate peripheral T cell survival in an IL-7 dependent mechanism.

### Sel1L/ERAD maintains naïve CD8^+^ T cell quiescence

In the absence of cognate antigen, peripheral T cell homeostasis is maintained through intrinsic and extrinsic mechanisms that promote cellular quiescence. After developing in the thymus, naïve (ie, antigen-inexperienced; CD44^lo^CD62L^hi^) T cells patrol the periphery in a quiescent state, with low proliferative, metabolic and translational rates. In addition to the survival defect noted in Sel1L-deficient peripheral T lymphocytes, we also noted a striking loss of the naïve CD8^+^ T cell population in the spleen and peripheral lymph nodes without a significant change in the naïve CD4^+^ T cell compartment (Figure 2A). Naïve T cell homeostasis is contingent on maintenance of cellular quiescence, which has classical hallmarks such as maintenance in the G0 phase of the cell cycle and low proliferative activity (Chapman et al., 2020). To examine if loss of the naïve CD8^+^ T cell compartment was due to loss of quiescence, we measured *in vivo* proliferation using Ki67, which is expressed in all phases of the cell cycle except G0, and by incorporation of the thymidine analog bromodeoxyuridine (BrdU) during S-phase over 24 hours. Sel1LKO naïve CD8^+^ T cells had increased BrdU incorporation (Figure 2B) and Ki67 expression (Figure 2C) compared to WT naïve cells, suggesting that Sel1L enforces naïve CD8^+^ T cell quiescence.

**Figure 2.**
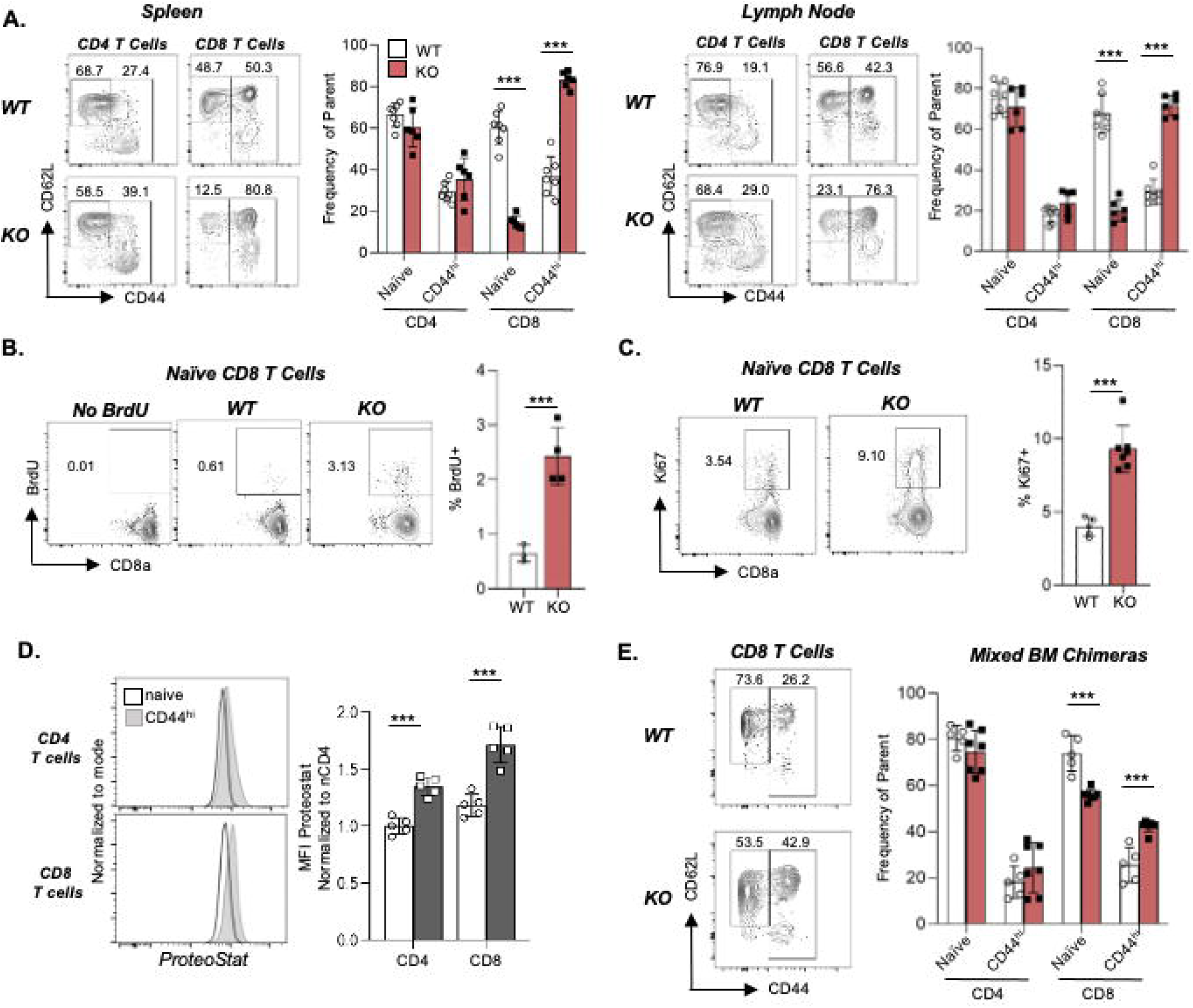
Sel1L maintains naïve CD8^+^ T cell homeostasis in a cell-intrinsic manner. Characterization of peripheral T cell subsets in Sel1LKO mice and littermate controls (WT), aged 6-10 weeks. A) *Left,* representative flow cytometric analysis of CD44 and CD62L expression on CD4^+^ and CD8^+^ T cells isolated from the spleens of WT and Sel1LKO mice. *Right,* frequency of naïve (CD44^lo^CD62L^hi^) and CD44^hi^ populations among CD4^+^ and CD8^+^ T cells isolated from the LN of WT and Sel1LKO mice. B) *Left,* representative flow cytometry of BrdU^+^ population in naïve CD8^+^ T cells; *right,* frequency of BrdU^+^ population in naïve CD8^+^ T cells in WT and Sel1LKO mice. C) *Left,* Ki67 expression in naïve CD8^+^ T cells from WT and Sel1LKO mice; *Right,* frequency of Ki67^+^ population among naïve CD8^+^ T cells in WT and Sel1LKO mice. D) *Left,* Proteostat staining in naïve and CD44^hi^ wild-type CD4^+^ and CD8^+^ T cells; *right,* MFI of Proteostat in indicated subsets, normalized to naïve CD4^+^ T cells. E) *Left,* representative flow cytometry of CD44 and CD62L in CD45.2^+^ WT or KO CD8^+^ T cells; *right,* frequency of naïve and CD44^hi^ subsets among CD8^+^ T cells derived from either WT or Sel1LKO BM in the mixed BM chimeras. Data are representative of n = 6-7/genotype (A), 3-4/genotype (B), 5-7/genotype (C, D), 5/genotype (E), from 2 (B-E) or 3 (A) independent experiments. Data are mean± SD. *p < 0.05, **p < 0.01, ***p < 0.001 (unpaired t-test). See also Supplementary Figure 2.

To determine if ERAD was active in wild-type T cells, we measured the level of protein aggresomes in naïve and CD44^hi^ (ie, activated or effector/memory) T cell subsets from wild-type mice using Proteostat. Both naïve CD4^+^ and CD8^+^ T cells had lower levels of protein aggresomes than CD44^hi^ subsets (Figure 2D), suggesting that ERAD may be active in naïve T cells.

T cell quiescence is enforced via intrinsic and extrinsic mechanisms (Chapman and Chi, 2018). Given that a lymphopenic environment can promote homeostatic proliferation and T cell activation (Goldrath et al., 2000), we asked if loss of CD8^+^ T cell quiescence is cell-intrinsic. Using the mixed bone marrow chimeras generated above, we found that CD8^+^ T cells derived from Sel1L-deficient donors had a smaller naïve compartment than CD8^+^ T cells derived from WT donors (Figure 2E) despite a non-lymphopenic environment. Together, these data confirm that Sel1L/ERAD preserves naïve CD8^+^ T cell homeostasis in a cell-intrinsic manner.

### Sel1L loss promotes mTORC1 activation and alters T cell metabolism

Regulation of intracellular metabolism, particularly through the mammalian target of rapamycin (mTOR) pathway, is critical to maintain T cell homeostasis. Loss of tuberous sclerosis complex 1 (Tsc1) results mTORC1 activation, diminished T cell survival and loss of quiescence (O’Brien et al., 2011; Pollizzi et al., 2015; Wu et al., 2011; Yang et al., 2011), which is a similar phenotype to Sel1LKO mice. Thus, we asked if Sel1L loss alters mTORC1 signaling. Sel1LKO naïve and CD44^hi^ CD8^+^ T cells had higher expression of the mTORC1 targets phosphorylated S6 (pS6; Figure 3A) and phosphorylated eukaryotic translation initiation factor 4E-binding protein (p4EBP1; Figure 3B) compared to control CD8^+^ T cells. There were no differences in pS6 or p4EBP1 expression in non-T lymphocytes in the WT or Sel1LKO mice (data not shown). Importantly, the activation of mTORC1 was also cell-intrinsic as evidenced by increased p4EBP1 in Sel1LKO T cells compared to competitor cells in the mixed BM chimeras (Figure 3C).

**Figure 3.**
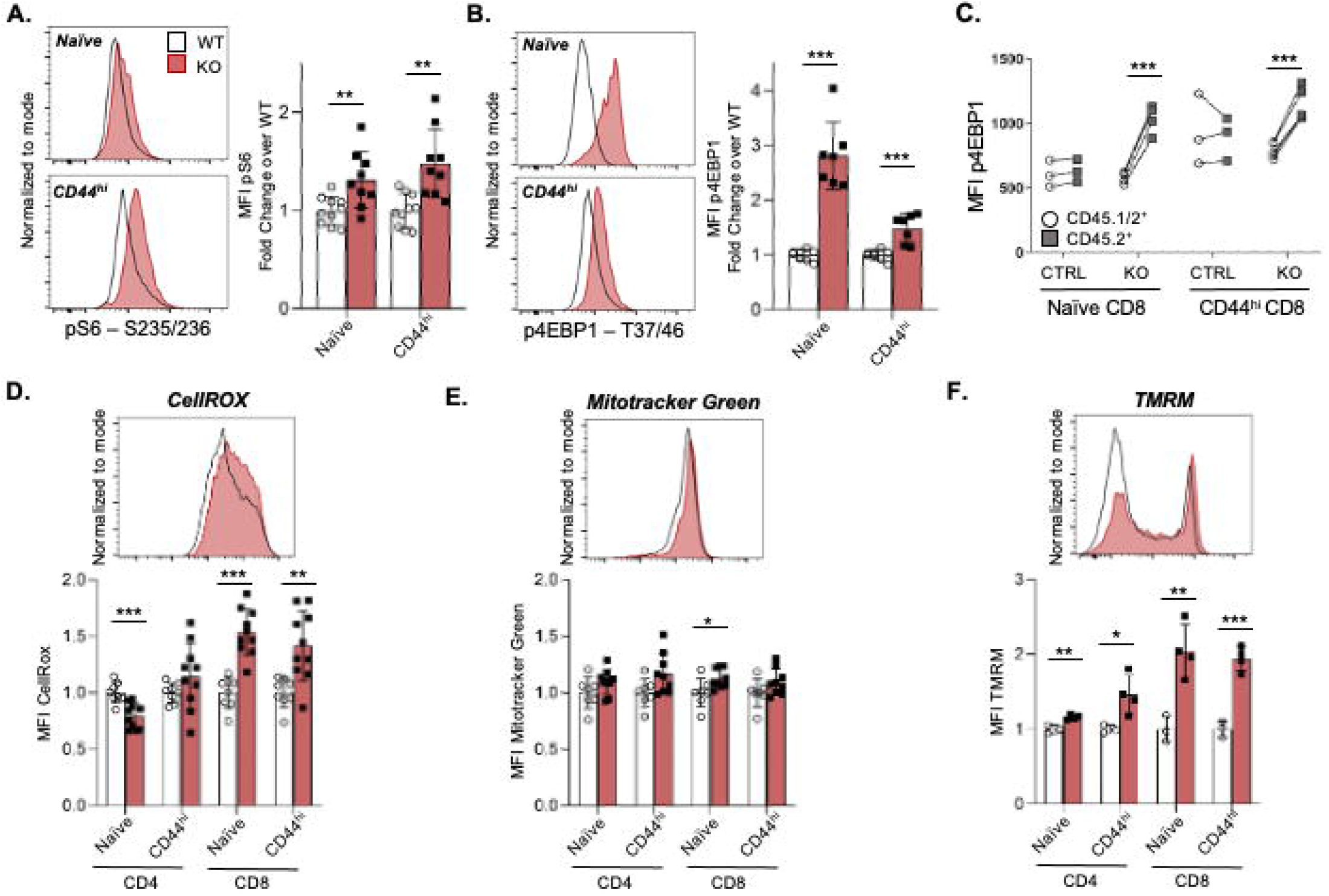
Sel1L Joss promotes mTORC1 activation and alters T ccU metabolism. Examination of mTORC1 targets and metabolic parameters in \1/Tand Sel1LKO T cells. A) *Left,* representative flow cytometric analysis of intracellular pS6 expression in indicated WT and Sel1LKO CD8^+^ T subsets; *right,* fold change in pS6 expression in naïve and CD44^hi^ CD8^+^ T cells from WT and KO mice, normalized to WT in each experiment. B) *Left,* representative flow cytometric analysis of intracellular p4EBP1 expression in indicated WT and Sel1LKO CDS8^+^ T subsets; *right,* fold change in p4EBP1 expression in naïve and CD44^hi^ CD8^+^ T cells from WT and Sel1LKO mice, normalized to WT in each experiment. C) MFI of p4EBP1 in splenic naïve and CD44hi CD8^+^ T cell subsets derived from competitor (CD45.1^+^CD45.2^+^) or experimental (WT or KO, CD45.2^+^) bone marrow in mixed bone marrow chimera mice. D-F) *Top,* representative flow cytometric plot of indicated fluorescent probe in WT and Sel1LKO naïve CD8^+^ T cells; *bottom,* MFI of indicated probe in the indicated T cell subsets in WT and Sel1LKO mice, normalized to WT in the individual experiments. Data representative of n=8-10/genotype (A, D), 7-9/genotype (B, E), 3-5/genotype (C, F), from 2 (C), 3 (B, D-F) or 4 (A) independent experiments. Data are mean± SD. *p < 0.05, **p < 0.01, ***p < 0.001 (unpaired t-test: A-B, D-F; paired t-test: C).

Since mTORC1 activation leads to altered cellular metabolism, we assessed several metabolic parameters in the Sel1L-deficient T cells. In T cells, Tsc1 deletion (and subsequent mTORC1 activation) leads to increased reactive oxygen species (ROS) (O’Brien et al., 2011; Yang et al., 2011) in both CD4^+^ and CD8^+^ T cells, as well as altered mitochondrial membrane potential (O’Brien et al., 2011; Wu et al., 2011; Yang et al., 2011). In the Sel1L-deficient peripheral T cells, we found increased cellular ROS in the naïve and CD44^hi^ CD8^+^ T cell subsets (Figure 3D), but not in the CD4^+^ T cell subsets. Additionally, Sel1LKO naïve CD8^+^ T cells had increased mitochondrial mass (Figure 3E) and mitochondrial membrane potential (Figure 3F). These data reveal that Sel1L loss alters cellular metabolism in T cells at steady state.

### Sel1L regulates CD8^+^ T cell immune function

We initially examined T cell receptor (TCR) induced responses in vitro to dissect the functional role of Sel1L in T cell activation. We activated WT and Sel1LKO T cells in vitro with αCD3/CD28 and first determined cell survival following TCR activation. We found that there was no difference in the frequency of live cells in the Sel1LKO T cells compared to WT T cells (Supplementary Figure 3A). Then we examined the upregulation of CD25 and CD69, two early markers downstream of TCR stimulation. Despite lower basal surface expression of TCRβ on the Sel1LKO CD8^+^ T cells (Supplementary Figure 3B), we found that similar frequencies of WT and Sel1L-deficient CD8^+^ T cells upregulated CD25 and CD69 after overnight stimulation (Supplementary Figure 3C). Sel1LKO CD8^+^ T cells expressed lower levels of CD69 on a per cell basis (Supplementary Figure 3D). When WT or Sel1LKO splenocytes were incubated in vitro with αCD3 for three days, both CD4 and CD8 T cells had proliferated to a similar extent (Supplementary Figure 3E). Together these data indicate that Sel1L is not required for TCR-induced activation and proliferation in vitro.

To investigate the role of Sel1L in T cell immune responses in vivo, we infected WT and Sel1LKO mice with recombinant *Listeria monocytogenes* engineered to express the model antigen LCMV glycoprotein 33-41 (LM-gp33). On days 7-8 post-infection, WT and Sel1LKO mice displayed a similar frequency of antigen-specific (gp33^+^) CD8^+^ T cells (Figure 4A) though with an expected decrease in the absolute number of gp33^+^ T cells in the Sel1LKO mice (Figure 4B). Following ex vivo stimulation, Sel1L deficiency resulted in decreased ability of the CD8^+^ T cells to produce tumor necrosis factor alpha (TNFα; Figure 4C) and IL-2 (Figure 4D) compared to control in response to both gp33-41 peptide and PMA/ionomycin stimulation. There were no differences in IFNγ production (data not shown). Additionally, Sel1L-deficient CD8^+^ T cells had lower poly-functionality (ie, production of multiple cytokines) compared to control CD8^+^ T cells (Figure 4E). Moreover, whereas WT mice cleared *Listeria* by day 7-8 post-infection, the majority of Sel1LKO mice could not (Figure 4F). Together these data demonstrate that Sel1L is critical for CD8^+^ T cell function in vivo.

**Figure 4.**
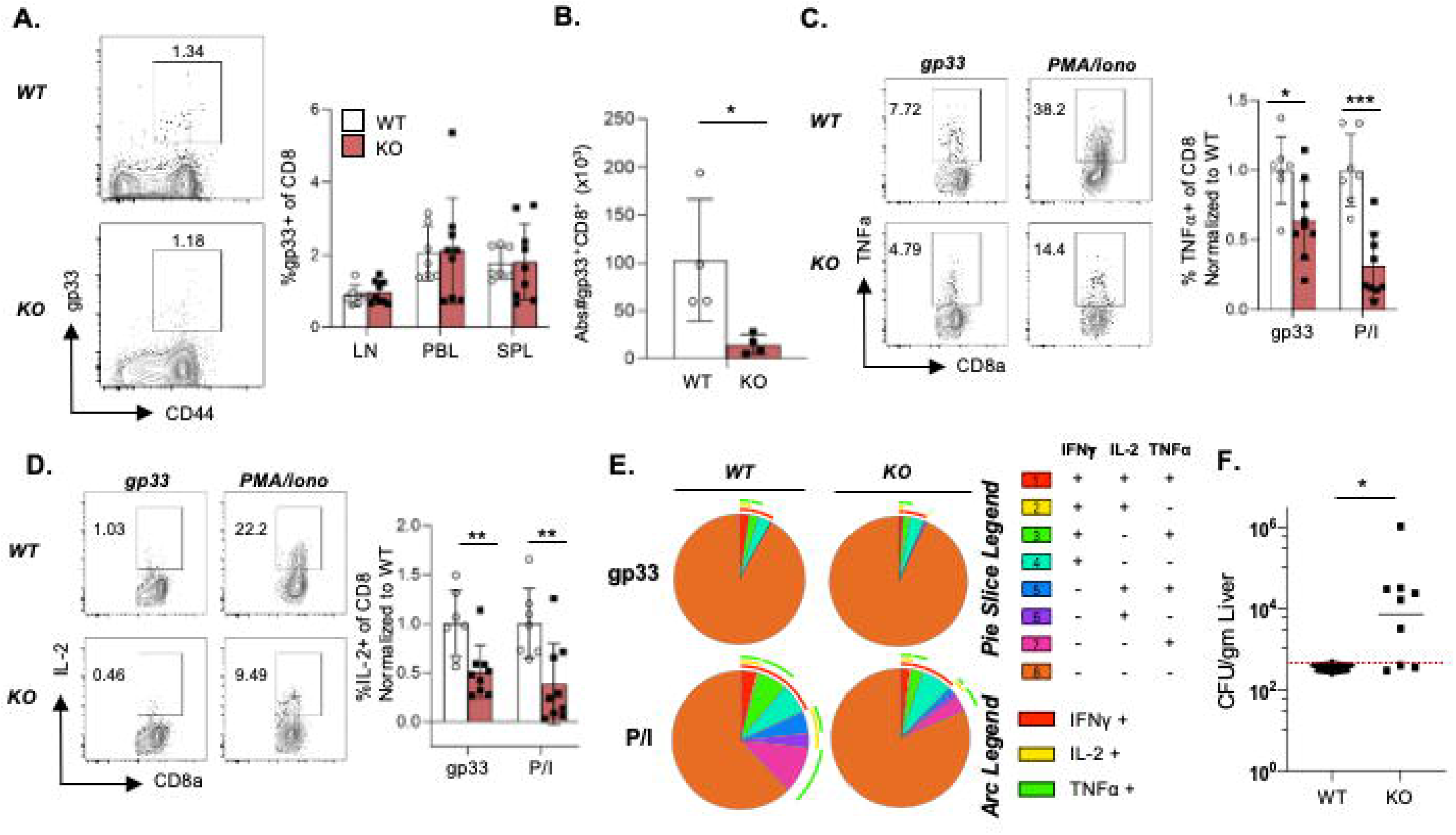
Sel1L regulates CD8^+^ T cell immune function. WT and Sel1LKO mice were infected with 5.8-9 x 10^4^ p.f.u. of *Listeria monocytogenes* expressing the gp33 epitope (LM-gp33). T cell responses were assessed on day 7-8 post-infection (p.i.). A) *Left,* representative flow cytometry of Gp33+ CD8+ T cells from WT and Sel1LKO splenocytes; *right,* frequency of gp33^+^ population among CD8^+^ T cells in the lymph nodes (LN), peripheral blood lymphocytes (PBLs) and spleen of WT and Sel1LKO mice. B) Absolute number of gp33^+^ CD8^+^ T cells in the spleen of WT and Sel1LKO mice. C-D) *Left,* representative flow cytometric analysis of intracellular TNFa (C) or IL-2 (D) in WT and Sel1LKO CD8^+^ T cells following gp33 and PMA/ionomycin (P/I) ex vivo stimulation; *right,* Frequency of TNFa^+^ or IL-2^+^ population among WT and Sel1LKO CD8^+^ T cells, normalized to WT, under indicated conditions. E) SPICE analysis of cytokine polyfunctionality an1ong WT and Sel1LKO CD8^+^T cells. F) Colony-fanning units (cfu) per gram of liver isolated from WT and KO mice, limit of detection represented by dashed line. Data representative of n=7-9/genotype, 2 independent experiments. Data are mean± SD. *p < 0.05, **p < 0.01, ***p < 0.001 (unpaired t test). See also Supplementary Figure 3.

## Discussion

In the present study, we identify Sel1L/ERAD as a critical factor that controls T cell survival, homeostasis and immune responses. Loss of Sel1L results in a mild reduction of single-positive thymocytes but a substantial loss of peripheral T cells with an increase in apoptotic cell death. Moreover, Sel1L/ERAD is required to maintain naïve CD8^+^ T cell homeostasis with Sel1L deletion leading to loss of naïve CD8^+^ T cell quiescence in a cell-intrinsic manner. Mechanistically, Sel1L loss leads to activation of mTORC1 signaling and altered cellular metabolism in the CD8^+^ T cell compartment, including increased cellular reactive oxygen species, mitochondrial membrane potential and mitochondrial mass. Furthermore, Sel1L is required for maximal CD8^+^ T cell immune responses following bacterial infection *in vivo*, and Sel1L deficiency leads to an impairment in cytokine production, polyfunctionality and pathogen clearance. These findings highlight the critical importance of Sel1L/ERAD in T lymphocyte homeostasis and function.

T cell specific deletion of either Se1lL, as detailed herein, or its binding partner the E3 ubiquitin ligase Hrd1 (Xu et al., 2016) results in profound peripheral T cell survival defect which supports a critical constitutive role for ERAD in T cell homeostasis. ERAD loss and accumulation of misfolded proteins can lead to ER stress and activation of the unfolded protein response (UPR) pathway. Prolonged intensity or duration of ER stress which the UPR cannot resolve will eventually trigger apoptosis (Hetz, 2012), as seen in Sel1L-deficient T cells. In contrast, loss of UPR mediators, including XBP1, IRE1α, CHOP (Cao et al., 2019; Govindarajan et al., 2018; Song et al., 2018), do not affect conventional T cell homeostasis at steady state suggesting that Sel1L/Hrd1-mediated ERAD but not UPR is more active and essential for T cell survival under physiologic conditions.

In addition to the survival defects, Sel1L also regulates naïve CD8^+^ T cell quiescence in a cell-intrinsic manner. These data are somewhat surprising on face value as naïve T cells have overall low protein synthesis rate compared to activated T cells (Araki et al., 2017; Kleijn and Proud, 2002), thus it would be predicted that there may be less need for ERAD and protein quality control in this quiescent subset. However, recent data have highlighted that even quiescent naïve T cells synthesize a total of ~60,000 proteins per minute (Wolf et al., 2020). Since translation is the most error-prone step in gene expression with approximately 10-30% of newly synthesized proteins undergoing ubiquitin-mediated degradation (Lykke-Andersen and Bennett, 2014; Schubert et al., 2000), even naïve T cells must require protein quality control systems. Tight control of messenger RNA (mRNA) levels by BTG1/BTG2, which promote mRNA deadenylation and degradation, and thus subsequent cellular protein levels, was recently identified as critical in the active maintenance of T cell quiescence (Hwang et al., 2020), supporting the hypothesis that proteostasis is critical to maintain T cell quiescence. Indeed, Sel1L has been recently identified as important to maintain cellular quiescence in hematopoietic stem cells (Liu et al., 2020; Xu et al., 2020) which raises the question if ERAD-mediated cellular proteostasis could be required to maintain quiescence in other cell types.

Further work needs to be done to fully dissect the mechanism by which Sel1L enforces T cell quiescence. One mechanism by which Sel1L may enforce T cell quiescence may be via repression of mTORC1 signaling in T cells as it has been well demonstrated that mTORC1 activity must be suppressed to maintain T cell quiescence (O’Brien et al., 2011; Wu et al., 2011; Yang et al., 2011). However, other work has also identified chronic ER stress mediated by a Schaflen 2 (Slfn2) mutation and subsequent UPR activation leading to loss of T cell quiescence (Omar et al., 2016). Additionally, a dysfunctional KDEL receptor 1 (KDELR1), an ER chaperone protein, leads to loss of naïve T cells through enhanced phosphorylation of eIF2α (Kamimura et al., 2015), a key downstream signaling molecule in the ER stress response. These findings raise the question if the ER stress triggered by Sel1L loss contributes to the loss of T cell quiescence. Complicating these studies is the well-identified UPR-mTORC cellular crosstalk. Chronic mTORC1 activation, such as following Tsc1/2 loss or in the Sel1L-deficient T cells, can promote ER stress-induced apoptosis (Bachar et al., 2009; Kato et al., 2012; Ozcan et al., 2008). Inhibition of mTORC1 signaling by rapamycin or protein synthesis by cycloheximide suppresses ER stress (Ozcan et al., 2008), suggesting that increased protein translation in Tsc-deficient cells may play a role in ER stress. Together these data do suggest that proteostasis may be an important cellular mechanism to maintain T cell quiescence.

In addition to regulating T cell homeostasis, we also demonstrate that Sel1L regulates CD8^+^ T cell immune responses. Other regulators of proteostasis, specifically those involved in the unfolded protein response (UPR), have also been shown to be critical in CD8^+^ T cell immunity and have been identified as potential novel targets for enhancing T cell immune responses. Deletion of XBP1 in T cells improved T cell mediated anti-tumor immunity (Song et al., 2018). Additionally, C/EBP homologous protein (CHOP) deletion enhances CD8^+^ T cell tumor immunity and improves CD8^+^ T cell responses following PD1 blockade (Cao et al., 2019). However, little is known about how ERAD regulates CD8^+^ T cell immunity. Our data show that Sel1L is critical for CD8^+^ T cell functional responses to *Listeria* infection with diminished IL-2 and TNFα production, as well as decreased bacterial clearance. Despite the profound survival defect at baseline, Sel1L-deficient CD8^+^ T cells do not show increased cell death following TCR stimulation and develop a similar proportion of antigen-specific T cells following infection. This finding was surprising given the significant increase in translational activity and protein production in activated T cells (Araki et al., 2017), which would be expected to have a concomitant increase in protein misfolding and ERAD load. UPR pathways are activated following TCR stimulation (Kemp et al., 2013; Thaxton et al., 2017), possibly compensating for any additional ER stress after T cell activation. The profound effect that T cell loss of Sel1L has on repressing pathogen clearance may be related to the decreased number of antigen-specific cells or could also be attributed to the significant defect in cytokine production among the existing CD8^+^ T cells. Further work is needed to dissect the respective cell-intrinsic roles of Sel1L in CD8^+^ T cell immunity.

## Acknowledgements

We thank Ling Qi for the Sel1L floxed mice. This work was funded through the National Institutes of Health by R01 HL150707 (QL), T32 AI007413 (LOC.), T32 CA140044 (LOC.), University of Michigan Rogel Cancer Center (SAC), Sheldon Glass Award (SAC), Limpert Scholar Award (SAC). Further support was provided by the National Cancer Institute of the National Institutes of Health (P30CA046592) by the use of the following Cancer Center Shared Resources: Flow Cytometry.

## Author contributions

ATD designed and performed experiments, as well as contributed to the writing/editing of the manuscript. LL, JPG, LOC and PK performed experiments. QL helped design experiments and provide conceptual insights. SAC designed experiments, performed experiments, wrote the manuscript and provided overall direction. All authors read and edited the manuscript.

## Methods

### Mice

C57BL/6J mice, and CD4Cre^+^ mice were obtained from The Jackson Laboratory and bred at the University of Michigan. B6.SJL-Ptprca (CD45.1^+^) mice were obtained from The Jackson Laboratory. Sel1L floxed mice (Sun et al., 2014) were a kind gift from Ling Qi. All mice were backcrossed at least ten times to a C57BL/6J background. Control mice for experiments included age-matched Sel1L^fl/fl^, Sel1L^+/+^ CD4Cre^+^, Sel1L^fl/fl^CD4Cre^-^ or C57BL/6J animals. All experiments were performed according to protocols approved by the Institutional Animal Care and Use Committee of the University of Michigan (PRO00009175; PRO00009775).

### Cell preparation and flow cytometry

Primary cell suspensions were obtained from thymi, spleens and lymph nodes. Cells were isolated, washed with FACS buffer (2% FBS in PBS), and stained with the indicated fluorophore conjugated anti-mouse antibodies, details of which are provided in the key resources table. Intracellular staining was performed using a Cytofix/Cytoperm kit (BD Biosciences) or a Foxp3 Transcription Factor Staining Buffer Set (eBioscience), according to the manufacturer’s instructions. Discrimination of live cell populations was performed using LIVE/DEAD Aqua or FarRed stains (Thermo Fischer) according to the manufacturer’s instructions prior to surface stainig. For Annexin V staining, cells were first surface stained for 30 minutes at 4C with surface antibodies, then washed with AnnexinV binding buffer and incubated with APC Annexin V as per manufacturer’s instructions (BioLegend). For phospho-flow cytometric staining, cells were fixed, permeabilized and stained with BD PhosphoFix (BD Biosciences) as per manufacturer’s instructions using pS6 or p4EBP1 primary antibodies (Cell Signaling) followed by AF488 goat anti-rabbit secondary (Invitrogen). For Proteostat staining, cells were stained with L/D viability dye and surface stained as previously indicated, then fixed with BD Cytofix/Cytoperm and stained with PROTEOSTAT Detection Reagent (Enzo Life Sciences). Fluorochrome-conjugated tetramers recognizing H2-D^b^–restricted gp33–41 epitope of LCMV were obtained from the National Institutes of Health Tetramer Core Facility. Cell sorting experiments were performed via Sony MA900 cell sorter. For flow cytometry experiments, data were acquired using BD Fortessa (BD Biosciences) and analyzed with FlowJo software (BD).

### Bone Marrow Chimeras

B6.SJL-Ptprca (CD45.1^+^) mice were irradiated with a total of 1100 rad in two divided doses and injected intravenously with a 1:1 mixture of T cell-depleted (magnetic bead depletion; Thy1.2; BioLegend) bone marrow from CD45.1^+^CD45.2^+^ mice and control C57BL/6J (CD45.2^+^) or Sel1L^fl/fl^CD4Cre^+^ (CD45.2^+^) mice. Recipient mice were maintained on antibiotic water for 4 weeks and then were analyzed 10–12 weeks after transplantation.

### BrdU labeling

Mice were injected intra-peritoneally with BrdU (Invitrogen) every 12hrs and given BrdU-containing water for 24 hours. After harvesting and processing organs, cells were stained as previously indicated then fixed with eBioscience Fix/Perm, treated with DNase for 60minutes and then intracellularly stained for 30 minutes with APC anti-BrdU antibody (Biolegend).

### In vitro T cell assays

Lymphocytes were isolated from spleen and lymph nodes, cultured in T cell media (10% FBS, 50 mM 2-ME, 2 mM L-glutamine/penicillin/streptomycin in IMDM) and activated with soluble αCD3 (2C11; eBioscience) at indicated concentrations and 5 mg/ml αCD28 (37.51; eBioscience) for the indicated times. For *in vitro* proliferation assay, cells were first labelled with CellTrace Violet (Fisher Scientific) as per manufacturer’s instructions, then cultured with soluble αCD3. Three days after activation, cells were surface stained with relevant flow cytometry antibodies prior to analysis. For cell survival assays, cells were incubated for indicated times with 10ng/mL of murine IL-7 (PeproTech) in complete media.

### Metabolism Assays

To measure metabolic parameters, 2×10^6^ splenocytes were incubated with 20uM MitoTracker Green in complete media, 500nM Tetramethylrhodamine Methyl Ester (TMRM) in PBS, or 2.5uM CellROX Green Reagent (all Invitrogen) for 30 minutes at 37°C prior to surface staining. For MitoTracker Green staining, 10mM verapamil (Sigma-Aldrich) was added to prevent dye efflux through xenobiotic efflux channels (Bonora et al., 2018; de Almeida et al., 2017).

### Real-time PCR

mRNA was isolated from indicated lymphocyte subpopulations using the RNAEasy Mini Kit (Qiagen) as per manufacturer’s instructions. RNA was quantified and converted into cDNA using SuperScript III First-Strand Synthesis System (Invitrogen). RT-PCR was performed using the primers listed in Key Resources Table. Relative gene expression of the target genes calculated using the 2^-ΔΔCT^ method and was normalized to β-actin expression levels.

### Western blotting

CD8^+^ T cells were negatively isolated via magnetic separation (Mojosort Mouse CD8+ T Cell Isolation Kit; BioLegend) and incubated with 10% Trichloric acid overnight at 4C. After centrifugation, protein pellets were washed with ice-cold acetone, air-dried and solubilized with solubilization buffer (9M Urea, 2% Triton X-100, 1% DTT) before NuPAGE LDS sample buffer was added. Proteins were separated on a NuPAGE 4-20% Tris-Glycine protein gels and transferred to a PVDF membrane.

### Infection

Mice were infected with 5.8-9.0×10^4^ CFU Listeria monocytogenes that expresses the LCMV gp33 epitope (LM-gp33), as indicated. *Ex vivo* stimulation of T cells was performed with 100ng/ml PMA and 1ug/ml ionomycin or 200mg/ml gp33 peptide (Anaspec) for 4-5 hours in the presence of brefeldin A. LM-gp33 was grown and bacterial loads were measured as previously described (Carty et al., 2018).

### Statistical analysis

Statistical significance was calculated as noted in the figure legends. Prism (GraphPad) was used for statistical analysis. SPICE (Simplified Presentation of Incredibly Complex Evaluations; NIH) was used for cytokine polyfunctionality analysis.

## References

Araki, K., Morita, M., Bederman, A.G., Konieczny, B.T., Kissick, H.T., Sonenberg, N., and Ahmed, R. (2017). Translation is actively regulated during the differentiation of CD8(+) effector T cells. Nat Immunol 18, 1046–1057.

Bachar, E., Ariav, Y., Ketzinel-Gilad, M., Cerasi, E., Kaiser, N., and Leibowitz, G. (2009). Glucose amplifies fatty acid-induced endoplasmic reticulum stress in pancreatic beta-cells via activation of mTORC1. PLoS One 4, e4954.

Bonora, M., Ito, K., Morganti, C., Pinton, P., and Ito, K. (2018). Membrane-potential compensation reveals mitochondrial volume expansion during HSC commitment. Exp Hematol 68, 30–37 e31.

Cao, Y., Trillo-Tinoco, J., Sierra, R.A., Anadon, C., Dai, W., Mohamed, E., Cen, L., Costich, T.L., Magliocco, A., Marchion, D., et al. (2019). ER stress-induced mediator C/EBP homologous protein thwarts effector T cell activity in tumors through T-bet repression. Nat Commun 10, 1280.

Carty, S.A., Gohil, M., Banks, L.B., Cotton, R.M., Johnson, M.E., Stelekati, E., Wells, A.D., Wherry, E.J., Koretzky, G.A., and Jordan, M.S. (2018). The Loss of TET2 Promotes CD8(+) T Cell Memory Differentiation. J Immunol 200, 82–91.

Chapman, N.M., Boothby, M.R., and Chi, H. (2020). Metabolic coordination of T cell quiescence and activation. Nat Rev Immunol 20, 55–70.

Chapman, N.M., and Chi, H. (2018). Hallmarks of T-cell Exit from Quiescence. Cancer Immunol Res 6, 502–508.

de Almeida, M.J., Luchsinger, L.L., Corrigan, D.J., Williams, L.J., and Snoeck, H.W. (2017). Dye-Independent Methods Reveal Elevated Mitochondrial Mass in Hematopoietic Stem Cells. Cell Stem Cell 21, 725–729 e724.

Goldrath, A.W., Bogatzki, L.Y., and Bevan, M.J. (2000). Naive T cells transiently acquire a memory-like phenotype during homeostasis-driven proliferation. J Exp Med 192, 557–564.

Govindarajan, S., Gaublomme, D., Van der Cruyssen, R., Verheugen, E., Van Gassen, S., Saeys, Y., Tavernier, S., Iwawaki, T., Bloch, Y., Savvides, S.N., et al. (2018). Stabilization of cytokine mRNAs in iNKT cells requires the serine-threonine kinase IRE1alpha. Nat Commun 9, 5340.

Hetz, C. (2012). The unfolded protein response: controlling cell fate decisions under ER stress and beyond. Nat Rev Mol Cell Biol 13, 89–102.

Jacobs, S.R., Michalek, R.D., and Rathmell, J.C. (2010). IL-7 is essential for homeostatic control of T cell metabolism in vivo. J Immunol 184, 3461–3469.

Johnston, J.A., Ward, C.L., and Kopito, R.R. (1998). Aggresomes: a cellular response to misfolded proteins. J Cell Biol 143, 1883–1898.

Kamimura, D., Katsunuma, K., Arima, Y., Atsumi, T., Jiang, J.J., Bando, H., Meng, J., Sabharwal, L., Stofkova, A., Nishikawa, N., et al. (2015). KDEL receptor 1 regulates T-cell homeostasis via PP1 that is a key phosphatase for ISR. Nat Commun 6, 7474.

Kato, H., Nakajima, S., Saito, Y., Takahashi, S., Katoh, R., and Kitamura, M. (2012). mTORC1 serves ER stress-triggered apoptosis via selective activation of the IRE1-JNK pathway. Cell Death Differ 19, 310–320.

Kemp, K.L., Lin, Z., Zhao, F., Gao, B., Song, J., Zhang, K., and Fang, D. (2013). The serine-threonine kinase inositol-requiring enzyme 1alpha (IRE1alpha) promotes IL-4 production in T helper cells. J Biol Chem 288, 33272–33282.

Kleijn, M., and Proud, C.G. (2002). The regulation of protein synthesis and translation factors by CD3 and CD28 in human primary T lymphocytes. BMC Biochem 3, 11.

Kong, S., Yang, Y., Xu, Y., Wang, Y., Zhang, Y., Melo-Cardenas, J., Xu, X., Gao, B., Thorp, E.B., Zhang, D.D., et al. (2016). Endoplasmic reticulum-resident E3 ubiquitin ligase Hrd1 controls B-cell immunity through degradation of the death receptor CD95/Fas. Proc Natl Acad Sci U S A 113, 10394–10399.

Liu, L., Inoki, A., Fan, K., Mao, F., Shi, G., Jin, X., Zhao, M., Ney, G., Jones, M.A., Sun, S., et al. (2020). ER associated degradation preserves hematopoietic stem cell quiescence and self-renewal by restricting mTOR activity. Blood.

Lykke-Andersen, J., and Bennett, E.J. (2014). Protecting the proteome: Eukaryotic cotranslational quality control pathways. J Cell Biol 204, 467–476.

O’Brien, T.F., Gorentla, B.K., Xie, D., Srivatsan, S., McLeod, I.X., He, Y.W., and Zhong, X.P. (2011). Regulation of T-cell survival and mitochondrial homeostasis by TSC1. Eur J Immunol 41, 3361–3370.

Omar, I., Lapenna, A., Cohen-Daniel, L., Tirosh, B., and Berger, M. (2016). Schlafen2 mutation unravels a role for chronic ER stress in the loss of T cell quiescence. Oncotarget 7, 39396–39407.

Ozcan, U., Ozcan, L., Yilmaz, E., Duvel, K., Sahin, M., Manning, B.D., and Hotamisligil, G.S. (2008). Loss of the tuberous sclerosis complex tumor suppressors triggers the unfolded protein response to regulate insulin signaling and apoptosis. Mol Cell 29, 541–551.

Pollizzi, K.N., Patel, C.H., Sun, I.H., Oh, M.H., Waickman, A.T., Wen, J., Delgoffe, G.M., and Powell, J.D. (2015). mTORC1 and mTORC2 selectively regulate CD8(+) T cell differentiation. J Clin Invest 125, 2090–2108.

Qi, L., Tsai, B., and Arvan, P. (2017). New Insights into the Physiological Role of Endoplasmic Reticulum-Associated Degradation. Trends Cell Biol 27, 430–440.

Rathmell, J.C., Farkash, E.A., Gao, W., and Thompson, C.B. (2001). IL-7 enhances the survival and maintains the size of naive T cells. J Immunol 167, 6869–6876.

Schubert, U., Anton, L.C., Gibbs, J., Norbury, C.C., Yewdell, J.W., and Bennink, J.R. (2000). Rapid degradation of a large fraction of newly synthesized proteins by proteasomes. Nature 404, 770–774.

Song, M., Sandoval, T.A., Chae, C.S., Chopra, S., Tan, C., Rutkowski, M.R., Raundhal, M., Chaurio, R.A., Payne, K.K., Konrad, C., et al. (2018). IRE1alpha-XBP1 controls T cell function in ovarian cancer by regulating mitochondrial activity. Nature 562, 423–428.

Sun, S., Shi, G., Han, X., Francisco, A.B., Ji, Y., Mendonca, N., Liu, X., Locasale, J.W., Simpson, K.W., Duhamel, G.E., et al. (2014). Sel1L is indispensable for mammalian endoplasmic reticulum-associated degradation, endoplasmic reticulum homeostasis, and survival. Proc Natl Acad Sci U S A 111, E582–591.

Thaxton, J.E., Wallace, C., Riesenberg, B., Zhang, Y., Paulos, C.M., Beeson, C.C., Liu, B., and Li, Z. (2017). Modulation of Endoplasmic Reticulum Stress Controls CD4(+) T-cell Activation and Antitumor Function. Cancer Immunol Res 5, 666–675.

Wolf, T., Jin, W., Zoppi, G., Vogel, I.A., Akhmedov, M., Bleck, C.K.E., Beltraminelli, T., Rieckmann, J.C., Ramirez, N.J., Benevento, M., et al. (2020). Dynamics in protein translation sustaining T cell preparedness. Nat Immunol 21, 927–937.

Wu, Q., Liu, Y., Chen, C., Ikenoue, T., Qiao, Y., Li, C.S., Li, W., Guan, K.L., Liu, Y., and Zheng, P. (2011). The tuberous sclerosis complex-mammalian target of rapamycin pathway maintains the quiescence and survival of naive T cells. J Immunol 187, 1106–1112.

Xu, L., Liu, X., Peng, F., Zhang, W., Zheng, L., Ding, Y., Gu, T., Lv, K., Wang, J., Ortinau, L., et al. (2020). Protein quality control through endoplasmic reticulum-associated degradation maintains haematopoietic stem cell identity and niche interactions. Nat Cell Biol 22, 1162–1169.

Xu, Y., Melo-Cardenas, J., Zhang, Y., Gau, I., Wei, J., Montauti, E., Zhang, Y., Gao, B., Jin, H., Sun, Z., et al. (2019). The E3 ligase Hrd1 stabilizes Tregs by antagonizing inflammatory cytokine-induced ER stress response. JCI Insight 4.

Xu, Y., Zhao, F., Qiu, Q., Chen, K., Wei, J., Kong, Q., Gao, B., Melo-Cardenas, J., Zhang, B., Zhang, J., et al. (2016). The ER membrane-anchored ubiquitin ligase Hrd1 is a positive regulator of T-cell immunity. Nat Commun 7, 12073.

Yang, K., Neale, G., Green, D.R., He, W., and Chi, H. (2011). The tumor suppressor Tsc1 enforces quiescence of naive T cells to promote immune homeostasis and function. Nat Immunol 12, 888–897.

