## Supplemental Figure 1 for "The endoplasmic reticulum associated degradation adaptor Sel1L regulates T cell homeostasis and function"

### Slide 1
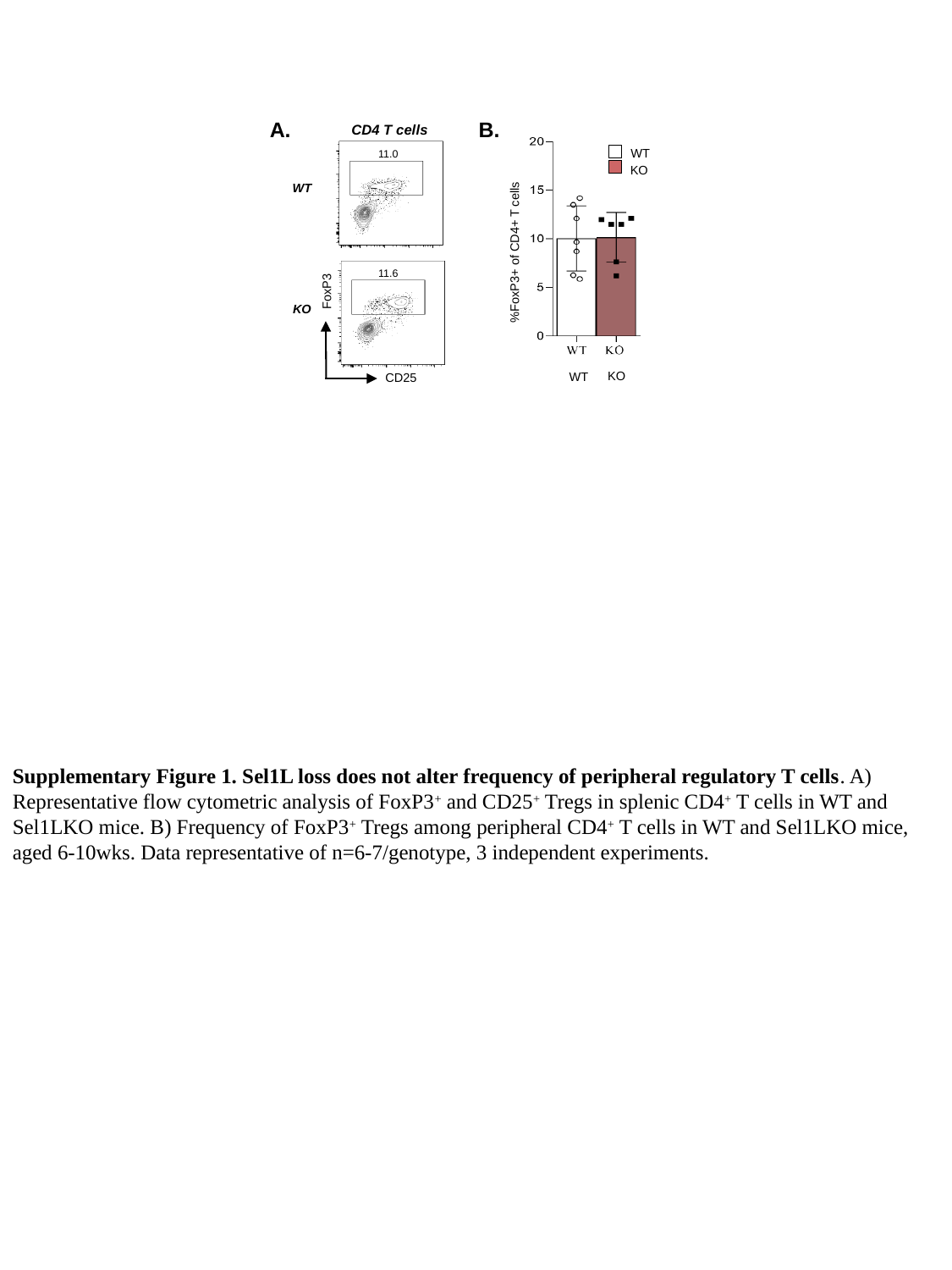

A.
KO
WT
%FoxP3+ of CD4+ T cells
11.0
WT
11.6
FoxP3
KO
CD25
B.
CD4 T cells
WT
KO
Supplementary Figure 1. Sel1L loss does not alter frequency of peripheral regulatory T cells. A) Representative flow cytometric analysis of FoxP3^+^ and CD25^+^ Tregs in splenic CD4^+^ T cells in WT and Sel1LKO mice. B) Frequency of FoxP3^+^ Tregs among peripheral CD4^+^ T cells in WT and Sel1LKO mice, aged 6-10wks. Data representative of n=6-7/genotype, 3 independent experiments.
