## Supplemental Figure 2 for "The endoplasmic reticulum associated degradation adaptor Sel1L regulates T cell homeostasis and function"

### Slide 1
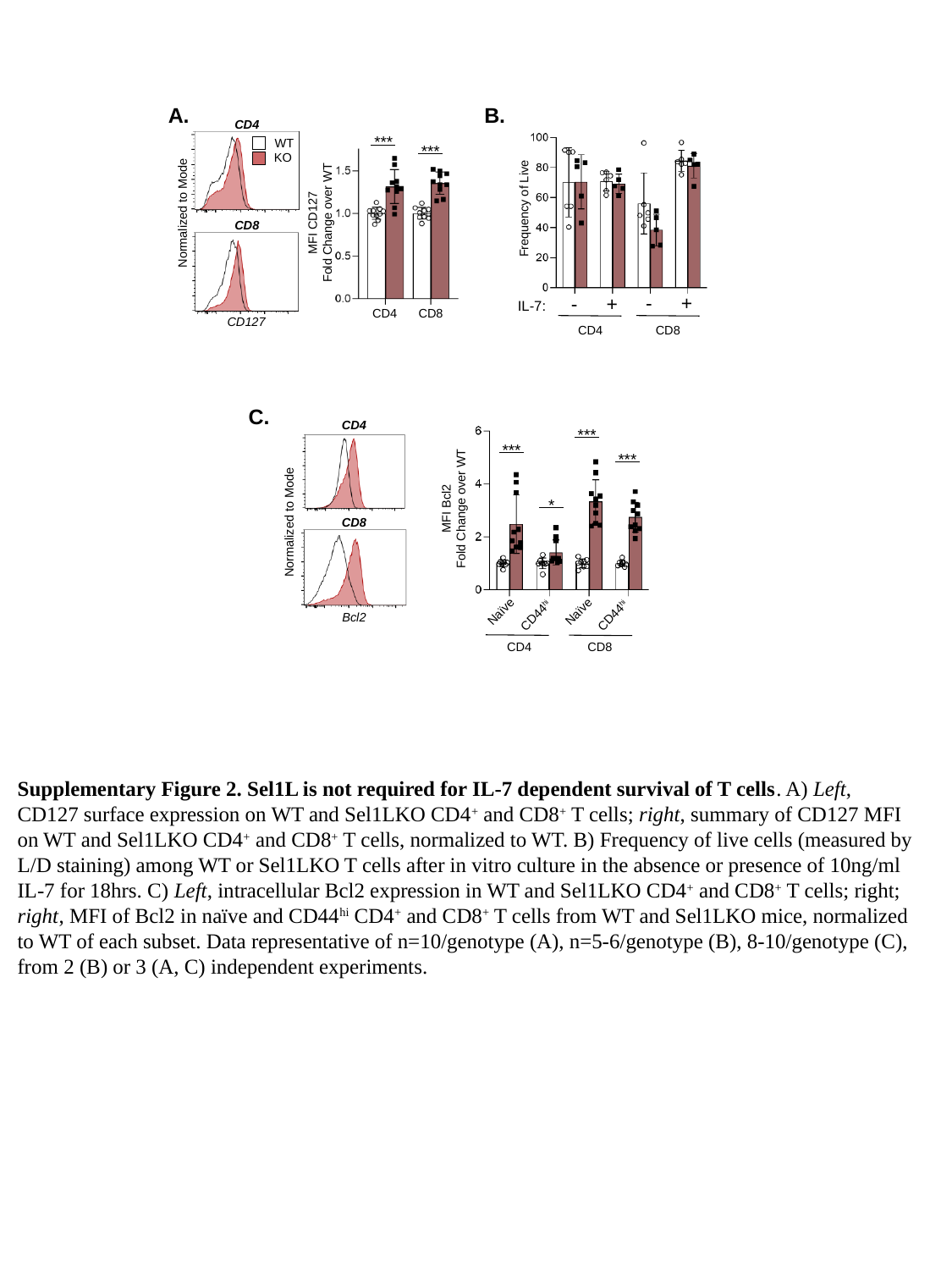

A.
B.
CD4
Normalized to Mode
CD8
CD127
***
CD4
CD8
MFI CD127
Fold Change over WT
***
Frequency of Live
CD4
CD8
IL-7:
WT
KO
-
+
-
+
***
***
***
*
Naïve
Naïve
CD44^hi^
CD44^hi^
CD4
CD8
MFI Bcl2
Fold Change over WT
C.
CD4
CD8
Normalized to Mode
Bcl2
Supplementary Figure 2. Sel1L is not required for IL-7 dependent survival of T cells. A) Left, CD127 surface expression on WT and Sel1LKO CD4^+^ and CD8^+^ T cells; right, summary of CD127 MFI on WT and Sel1LKO CD4^+^ and CD8^+^ T cells, normalized to WT. B) Frequency of live cells (measured by L/D staining) among WT or Sel1LKO T cells after in vitro culture in the absence or presence of 10ng/ml IL-7 for 18hrs. C) Left, intracellular Bcl2 expression in WT and Sel1LKO CD4^+^ and CD8^+^ T cells; right; right, MFI of Bcl2 in naïve and CD44^hi^ CD4^+^ and CD8^+^ T cells from WT and Sel1LKO mice, normalized to WT of each subset. Data representative of n=10/genotype (A), n=5-6/genotype (B), 8-10/genotype (C), from 2 (B) or 3 (A, C) independent experiments.
