## Supplemental Figure 3 for "The endoplasmic reticulum associated degradation adaptor Sel1L regulates T cell homeostasis and function"

### Slide 1
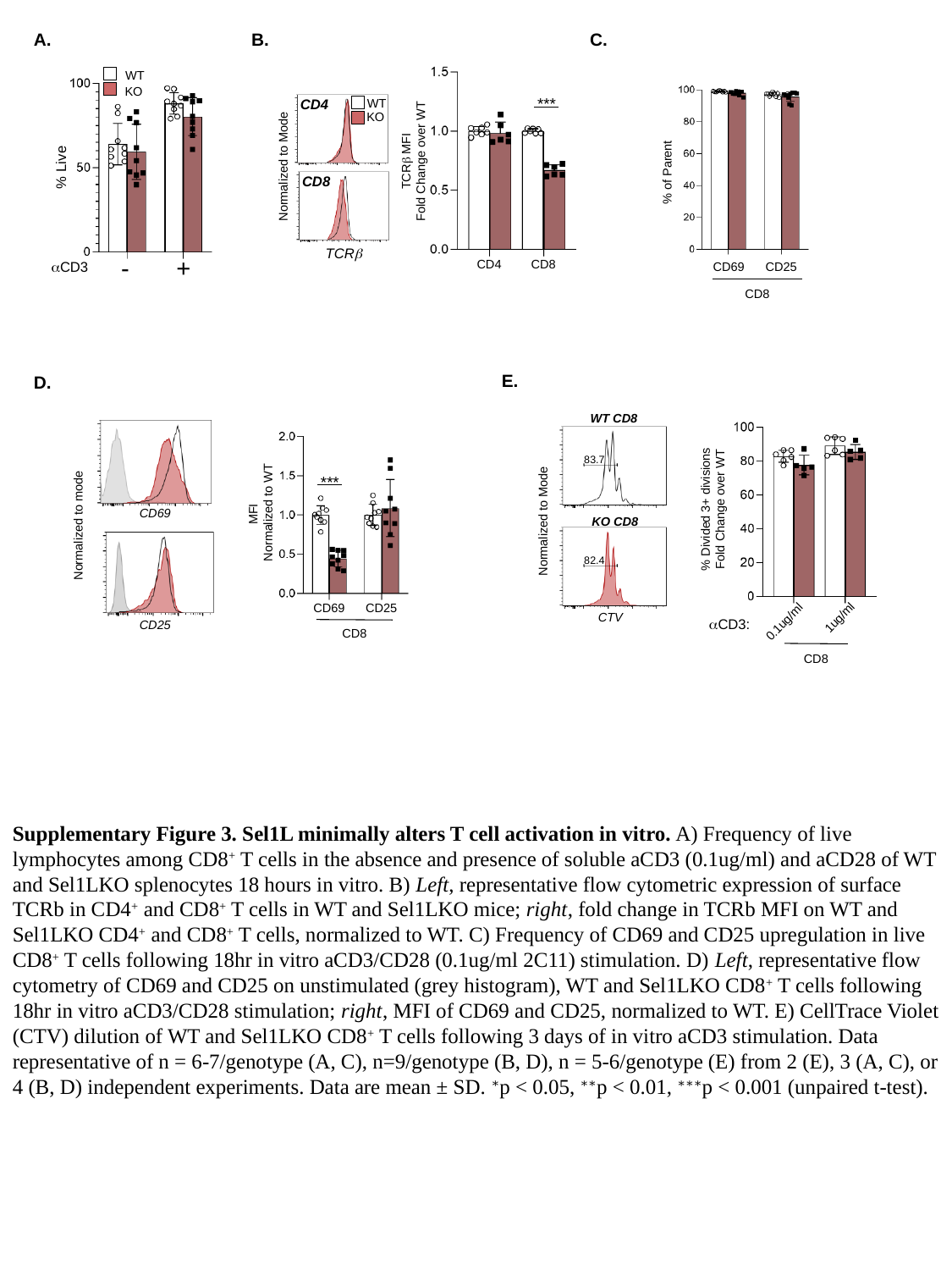

B.
A.
C.
CD4
Normalized to Mode
CD8
TCRβ
TCRβ MFI
Fold Change over WT
CD8
CD4
***
WT
KO
% Live
-
+
αCD3
WT
KO
% of Parent
CD69
CD25
CD8
E.
D.
MFI
 Normalized to WT
CD69
CD25
CD8
***
WT CD8
Normalized to Mode
CTV
83.7
82.4
KO CD8
% Divided 3+ divisions
Fold Change over WT
1ug/ml
0.1ug/ml
CD8
αCD3:
Normalized to mode
CD25
CD69
Supplementary Figure 3. Sel1L minimally alters T cell activation in vitro. A) Frequency of live lymphocytes among CD8^+^ T cells in the absence and presence of soluble aCD3 (0.1ug/ml) and aCD28 of WT and Sel1LKO splenocytes 18 hours in vitro. B) Left, representative flow cytometric expression of surface TCRb in CD4^+^ and CD8^+^ T cells in WT and Sel1LKO mice; right, fold change in TCRb MFI on WT and Sel1LKO CD4^+^ and CD8^+^ T cells, normalized to WT. C) Frequency of CD69 and CD25 upregulation in live CD8^+^ T cells following 18hr in vitro aCD3/CD28 (0.1ug/ml 2C11) stimulation. D) Left, representative flow cytometry of CD69 and CD25 on unstimulated (grey histogram), WT and Sel1LKO CD8^+^ T cells following 18hr in vitro aCD3/CD28 stimulation; right, MFI of CD69 and CD25, normalized to WT. E) CellTrace Violet (CTV) dilution of WT and Sel1LKO CD8^+^ T cells following 3 days of in vitro aCD3 stimulation. Data representative of n = 6-7/genotype (A, C), n=9/genotype (B, D), n = 5-6/genotype (E) from 2 (E), 3 (A, C), or 4 (B, D) independent experiments. Data are mean ± SD. ^∗^p < 0.05, ^∗∗^p < 0.01, ^∗∗∗^p < 0.001 (unpaired t-test).
